# Near-atomic cryo-EM structure of yeast kinesin-5-microtubule complex reveals a distinct binding footprint

**DOI:** 10.1101/302455

**Authors:** Ottilie von Loeffelholz, Alejandro Peña, Douglas Robert Drummond, Robert Cross, Carolyn Ann Moores

## Abstract

Kinesin-5s are essential members of the superfamily of microtubule-dependent motors that undertake conserved roles in cell division. We investigated coevolution of the motor-microtubule interface using cryo-electron microscopy to determine the near-atomic structure of the motor domain of Cut7, the fission yeast kinesin-5, bound to fission yeast microtubules. AMPPNP-bound Cut7 adopts a kinesin-conserved ATP-like conformation, with a closed nucleotide binding pocket and docked neck linker that supports cover neck bundle formation. Compared to mammalian tubulin microtubules, Cut7’s footprint on *S. pombe* microtubule surface is subtly different because of their different architecture. However, the core motor-microtubule interaction that stimulates motor ATPase is tightly conserved, reflected in similar Cut7 ATPase activities on each microtubule type. The *S. pombe* microtubules were bound by the drug epothilone, which is visible in the taxane binding pocket. Stabilization of *S. pombe* microtubules is mediated by drug binding at this conserved site despite their noncanonical architecture and mechanochemistry.

**Highlights:**

- *S. pombe* Cut7 has a distinct binding footprint on *S. pombe* microtubules
- The core interface driving microtubule activation of motor ATPase is conserved
- The neck linker is docked in AMPPNP-bound Cut7 and the cover neck bundle is formed
- Epothilone binds at the taxane binding site to stabilize *S. pombe* microtubules

**eTOC text:** To investigate coevolution of the motor-microtubule interface, we used cryo-electron microscopy to determine the near-atomic structure of the motor domain of Cut7, the fission yeast kinesin-5, bound to microtubules polymerized from natively purified fission yeast tubulin and stabilised by the drug epothilone.

## Introduction

Members of the kinesin superfamily of microtubule-based, ATP-driven molecular motors play multiple essential roles in cell division, reflecting the dynamic complexity of the machinery required for accurate chromosome segregation. Kinesin-5 motors are conserved amongst eukaryotes and undertake related roles in cell division, driving spindle pole separation and stabilisation of spindle bipolarity (Goulet and Moores, 2013). These roles are enabled by the quaternary structure of kinesin-5s, with their tetrameric, dumbbell shape supporting spindle microtubule (MT) cross linking and sliding. Cut7, the sole kinesin-5 in fission yeast, was amongst the first mitotic kinesins to be identified and is essential for spindle pole body separation (Hagan and Yanagida, 1990). While most kinesin-5 motors take steps towards MT plus ends (Kapitein et al., 2005), several yeast kinesin-5s including Cut7, have been shown to move to MT plus or minus ends according to molecular context (Britto et al., 2016; Edamatsu, 2014; Fridman et al., 2013; Gerson-Gurwitz et al., 2011; Roostalu et al., 2011). The precise molecular mechanism(s) of direction change in these yeast motors are not known, but presumably the relatively small kinesin repertoire in these organisms (9 kinesins in the fission yeast genome, 6 in budding yeast) is one evolutionary driver for encoding regulated bidirectionality in single motors.

An important and coevolved facet of motor functionality is the motor-MT interface. αβ-tubulin heterodimers are the building blocks of MTs and are among the most highly conserved proteins in eukaryotes. However, the specificity of the kinesin-MT interaction suggests that even subtle differences in the MT track could influence motor function. Tubulin purified from mammalian brain remains the default substrate for investigation of kinesin function. However, it is a heterogeneous mix of multiple tubulin isoforms and post-translationally modified tubulins, so although these sources of diversity influence motor properties (Sirajuddin et al., 2014), their individual contributions to motor function cannot be discriminated. Recent advances in preparation of tubulin from diverse biological sources is highlighting the potential for diverse behavior even among nominally highly conserved tubulins (Drummond et al., 2011; Howes et al., 2017; Minoura et al., 2013; Uchimura et al., 2006; von Loeffelholz et al., 2017; Widlund et al., 2012). Purified native fission yeast tubulin contains a mixture of α-tubulin1/2 isoforms (87% sequence identity), a single β-tubulin isoform and no post-translational modifications (Drummond et al., 2011). We recently showed that the structural properties of fission yeast tubulin (abbreviated here to Sp_tub) are distinct from mammalian brain tubulin (Mam_tub), including polymerization of skewed protofilaments within the MT lattice and a different structural response to tubulin GTPase activity (von Loeffelholz et al., 2017). Furthermore, the sensitivity of yeast tubulin to MT-binding drugs differs from mammalian tubulin (Barnes et al., 1992; Bode et al., 2002).

Exploiting the availability of purified native fission yeast tubulin and recombinantly expressed Cut7 motor domain (Cut7MD), we determined the structure of this yeast motor bound to its native MT track at near-atomic resolution using cryo-electron microscopy (cryo-EM). This is the best-resolved reconstruction of a MT-bound kinesin-5 motor determined to date and allows us to map the motor-MT interface precisely. We use our structure to compare the footprint of this yeast motor on Sp_tub MTs with the previously determined Cut7 interaction with Mam_tub MTs (Britto et al., 2016), and we investigate the effect of these differences in the motor-track interface on steady-state motor ATPase activity. Finally, we visualize the binding site of the drug epothilone on Sp_tub MTs, shedding light on its MT stabilization mechanism and mode of influence on dynamic instability in yeast MTs.

## Results and Discussion

### Near-atomic structure of the ATP-like conformation of yeast Cut7 motor domain bound to yeast MTs

Low dose cryo-EM movies of Cut7MD in the presence of the non-hydrolyzable ATP analogue AMPPNP bound to Sp_tub MTs stabilized by the drug epothilone were collected. The regular, 8nm-spaced binding characteristic of most kinesin motor domains – e.g. (Atherton et al., 2014; Shang et al., 2014) - was also observed for Cut7MD (Figure S1A). This is distinct, however, from the super-stoichiometric binding observed for the budding yeast kinesin-5 Cin8 motor domain (Bell et al., 2017), which depends on a large insert in loop8 that is not present in Cut7. Thus, yeast kinesin-5 motor constructs can exhibit divergent properties, but in our cryo-EM experiment Cut7MD displays a canonical kinesin binding pattern every tubulin dimer on Sp_tub MTs.

Motion-corrected movies of Cut7MD-bound 13 PF Sp_tub MTs were processed using a previously described set of custom-written scripts (Sindelar and Downing, 2007; von Loeffelholz et al., 2017). The overall resolution of the Cut7MD-MT reconstruction is 4.5 Å (0.143 FSC criterion; Figure S1B, C). As is typical in motor-MT complexes (Kellogg et al., 2017), there is a resolution gradient from MT to kinesin density (Figure S1D). The average resolution in the tubulin region of the reconstruction is ~ 4.4 Å with some regions substantially better than this. The average resolution in the kinesin region of the reconstruction is 5.4 Å, and this density was sharpened independently of the MT region (Figure 1A). Using the cryo-EM density, pseudo-atomic models of *S. pombe* α-tubulin1, β-tubulin and Cut7MD were built.

**Figure 1.**
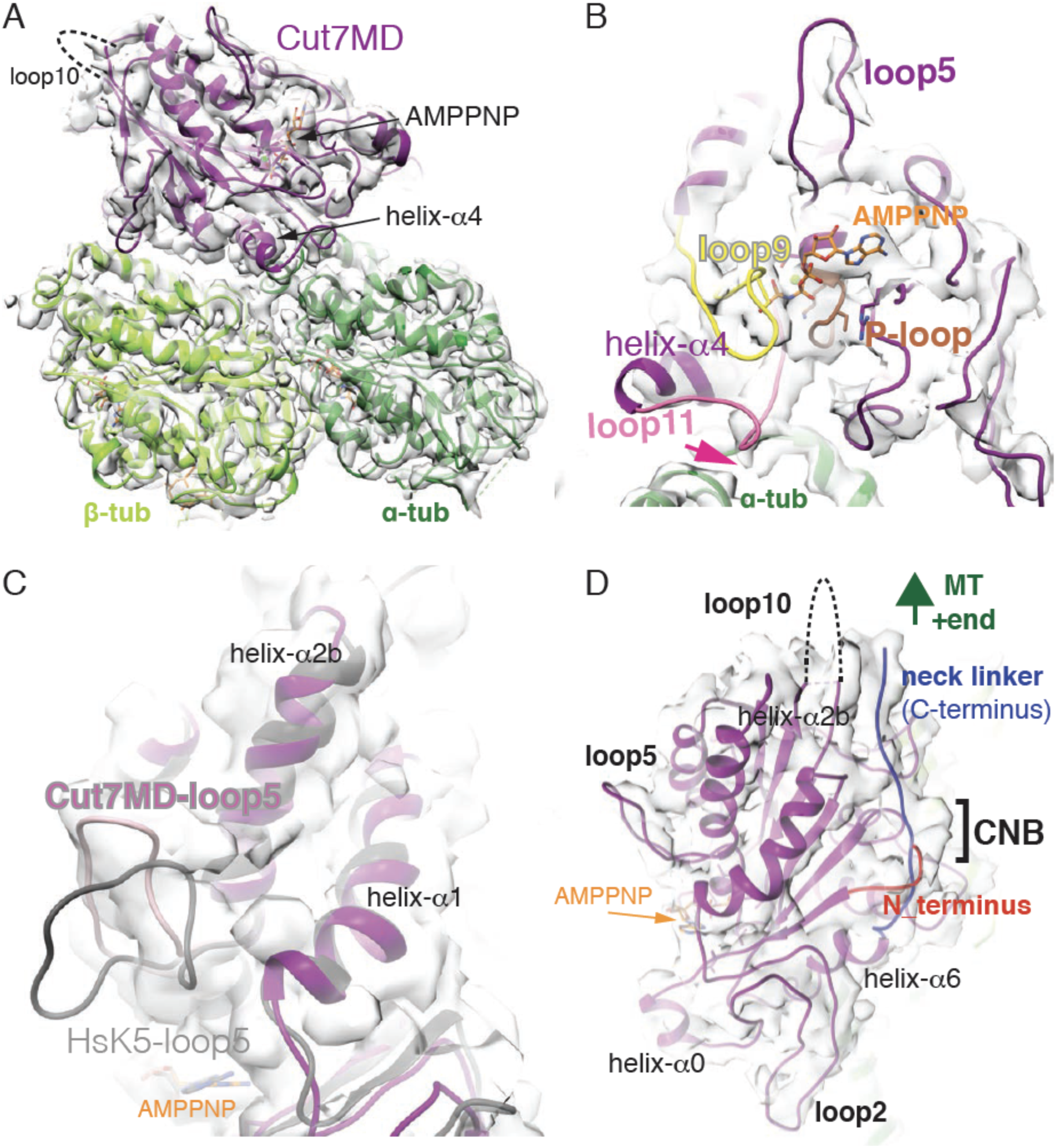
The near-atomic resolution reconstruction of *S. pombe* Cut7MD-AMPPNP bound to Sp_tub MT. (A) The asymmetric unit (αβ-tubulin + Cut7MD) of the reconstruction viewed towards the nucleotide binding pocket; the position of the disordered loop 10 is indicated by the dotted line; (B) View of the Cut7MD nucleotide binding pocket showing helix-α4 at the MT surface, the conserved nucleotide coordinating loops, P-loop (brown), loop9 (yellow) and loop11 (pink), as well as loop5 (purple) emerging away from the nucleotide binding site; the pink arrow indicates the separation of density corresponding to loop11 and the MT surface; (C) Close up view of the distinct conformation of Cut7MD loop5 (pink) compared to human kinesin-5 (HsK5-AMPPNP (PDB:3HQD), in grey); (D) Cut7MD cover neck bundle (CNB) formation from the docked neck linker (blue) – directed towards the MT plus end - and the N-terminus (red).

Cut7MD adopts the same orientation with respect to the Sp_tub MT lattice as previously seen on Mam_tub MTs (Britto et al., 2016) – albeit now visualized at substantially higher resolution – with helix-α4 of Cut7MD centred on the αβ-tubulin intradimer interface (Figure 1A). Consistent with sample preparation conditions, there is strong density in the nucleotide binding pocket corresponding to bound Mg-AMPPNP (Figure 1A, B). The conserved nucleotide-binding loops – the P-loop, loop 9 (containing switch I), loop 11 (containing switch II) (Figure S2) – adopt the compact conformation seen in other kinesin motors in the presence of this nucleotide analogue (Figure 1B) (Atherton et al., 2014; Britto et al., 2016; Gigant et al., 2013; Parke et al., 2010; Shang et al., 2014). The observation that the loop11 helical turn is not connected to the MT surface (Figure 1B, pink arrow) is also consistent with previously characterised ATP-like conformations.

At the plus end of the motor domain, there is no clear density visible corresponding to the 17 amino acid insert in loop10 (Figure 1A), suggesting that this loop is flexible and is thus not readily visualised at the high resolution of our current structure. However, immediately above and protruding away from the nucleotide binding site, density corresponding to the Cut7MD-specific loop5 is clearly visible (Figure 1B, C). As was previously seen at lower resolution (Britto et al., 2016), this Cut7MD loop5 conformation is distinct from that observed for AMPPNP-bound human kinesin-5 loop5 (Goulet et al., 2012; Parke et al., 2010). The sequences of Cut7 and human kinesin-5 in this region of the motor domain are distinct (Figure S2), and conformational sensitivity of this loop to even small changes in other kinesin-5s has been observed (Figure 1C; (Behnke-Parks et al., 2011; Bell et al., 2017; Bodey et al., 2009; Liu et al., 2011). Loop5 in human kinesin-5 forms part of a binding pocket for small molecule inhibitors (Goulet and Moores, 2013) and in addition to conformational divergence, several residues in human kinesin-5 that mediate inhibitor binding (e.g. W127, Y211) are not conserved in Cut7MD (Figure S2). Thus, our structure predicts that Cut7MD would be insensitive to human-specific kinesin-5 inhibitors, but small molecules designed to bind to this region of fungal kinesin-5s might be expected to be interesting candidates for fungicides.

Distal to the nucleotide binding site, the Cut7MD N- and C-termini coincide to form the functionally important cover neck bundle (CNB; Figure 1D) (Goulet et al., 2012; Khalil et al., 2008). Density corresponding to the C-terminal neck linker, which connects the Cut7 motor domain to the rest of the protein, is clearly visible docking along the length of the motor domain and directed towards the MT plus end. This docked conformation is consistent with the AMPPNP state of the motor and with canonical concepts of plus-end directed motor movement, which has been described for Cut7MD monomers (Britto et al., 2016; Rice et al., 1999). In this conformation, a short portion of the N-terminus lies above the docked neck linker, forming the CNB that contributes to force generation by kinesins.

### The Cut7-microtubule interaction

Cut7MD binds to Sp_tub MTs using canonical kinesin structural elements as were previously seen in the lower resolution reconstruction of the motor bound to Mam_tub MTs (Britto et al., 2016): helix-α4 at the intradimer interface, with additional contacts formed with β-tubulin (primarily H12) by helix-α5 and β5/loop8 and with α-tubulin (H12) by helix-α6 (Figure 2A; Figure S2). The high degree of sequence conservation between Sp_tub and Mam_tub means that the residues on the surface of each MT are essentially identical (Figure 2A, (von Loeffelholz et al., 2017)). However, our previous work also showed that the intrinsic architecture of *in vitro* polymerized Sp_tub MTs are different compared to Mam_tub MTs: PFs in Sp_tub MTs are more skewed and no nucleotide-dependent lattice compaction is observed (von Loeffelholz et al., 2017). As a consequence, the footprint of Cut7MD on Sp_tub MTs (Figure 2B) is subtly different compared with that on Mam_tub MTs (Figure 2C), being slightly skewed towards β-tubulin H3.

**Figure 2.**
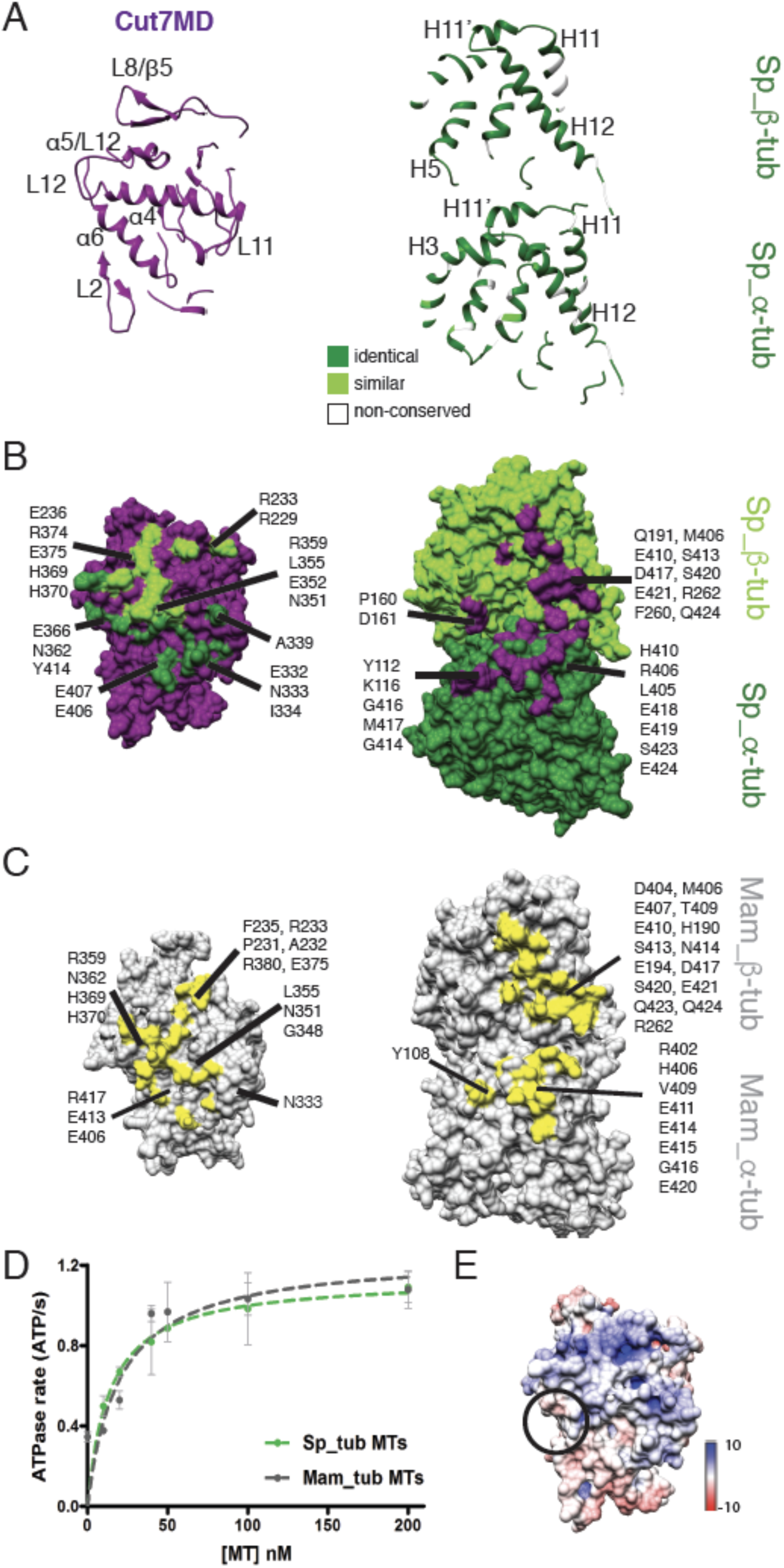
The Cut7 footprint on Sp_tub MTs is distinct. (A) Ribbon depiction of longitudinal slices through the Cut7MD model (left) and Sp_tub αβ-tubulin (right) showing the structural elements involved in binding between motor and MT track; the αβ-tubulin ribbon is colored according to sequence conservation with mammalian tubulin; note, while loop2 (L2) lies close to the MT surface, there is no evidence in the EM density of a direct connection between this loop and the MT (B) View of the MT binding surface of Cut7MD (left) and view of the Cut7MD binding surface of Sp_tub αβ-tubulin (right) with residues <4 Å distant from each binding partner labelled and colored green and purple, respectively; (C) View of the MT binding surface of Cut7MD (left) and view of the Cut7MD binding surface of Mam_tub αβ-tubulin (right) with residues <4 Å distant from each binding partner labelled and colored yellow; (D) Cut7MD steady-state ATPase rate plotted as a function of [MT] for Sp_tub (green) and Mam_tub (grey). Data were fit to a Michaelis Menten kinetic yielding values for Cut7MD V_max_ and K_0.5_MT respectively of 1.14 ATP/s and 14nM on Sp_tub MTs and 1.25 ATP/s and 19.9nM on Mam_tub MTs, differences which are not statistically significant (p>0.99). (E) Cut7MD MT binding interface colored by surface charge. The approximate position of the adjacent negatively charged β-tubulin C-terminal tail is marked by a circle.

To evaluate the effect of this MT configuration on motor activity, we measured the steady state ATPase activity of Cut7MD on Sp_tub MTs and Mam_tub MTs, which showed that the Cut7MD ATPase rates on each MT type are not statistically significant different (V_max_ = 1.14 ATP/s on Sp_tub MTs and V_max_ = 1.25 ATP/s on Mam_tub MTs, Figure 2D). Characteristic of kinesin-5s (Goulet and Moores, 2013), Cut7MD has a very slow ATPase *in vitro*, which may be important in the context of its spindle function.

Point mutagenesis has revealed the role of specific residues at the kinesin-MT interface in stimulating kinesin ATPase and thus function e.g. (Uchimura et al., 2006). Our work provides a different view of the motor-MT interface, in which the underlying architecture of the MT slightly alters otherwise very similar binding interfaces. Despite evidence that kinesin motor activity is sensitive to the stabilization mode of their underlying MT tracks e.g. (Vale et al., 1994), we find that alterations in the binding interface resulting from differences in the shape of the tubulin dimer and the lattice architecture can be accommodated with no measurable alteration in kinesin ATPase.

Although the differences in Cut7MD footprint on each MT type are not evident in the ATPase activity of the monomer, they are likely to manifest in the stepping properties of larger constructs - including the emergent property of direction reversibility – and MT cross-linking activity of full length Cut7 tetramers (Britto et al., 2016). Our cryo-EM reconstructions do not provide structural information about another crucial point of difference between these two tubulin sources: the highly flexible, and therefore structurally invisible tubulin C-terminal tails. In mammalian brain tubulin, the C-terminal tails of α- and β-tubulin are variable between isoforms and subject to multiple post-translational modifications, e.g. (Sirajuddin et al.,
2014), whereas the Sp_tub has none (Drummond et al., 2011). Interestingly, the region of Cut7MD closest to the β-tubulin C-terminal tail is somewhat apolar (Figure 2E), perhaps helping to explain the absence of apparent interaction between the motor and MT at this site. However, the C-terminal tails of both tubulins could influence the activity of full length Cut7. Future experiments will further explore the mechanistic evidence for such effects.

### Visualization of epothilone bound to *S. pombe* MTs

The Sp_tub MTs in our reconstruction were stabilized by the drug epothilone. Epothilone has previously been shown to bind to the taxane pocket of β-tubulin that faces the MT lumen (Prota et al., 2013) and thereby stabilize Mam_tub and tubulin from budding yeast (Sc_tub) (Bode et al., 2002; Howes et al., 2017; Prota et al., 2013). In our reconstruction, density corresponding to epothilone in this binding site is clear (Figure 3A, B), demonstrating that this drug binds to Sp_tub in a similar way, despite some sequence divergence in the M-loop region (e.g. Sp_tub S279; Mam_tub Q281). Comparison with our previous structure of Sp_tub MTs bound by the protein Mal3, in which the taxane pocket is empty (Figure 3B) (von Loeffelholz et al., 2017), reinforces the assignment of this density. Although the resolution of our reconstruction is not sufficient to allow *de novo* determination of the configuration of the bound drug, our density is consistent with the epothilone orientation revealed by X-ray crystallography in complex with Mam_tub dimers (Prota et al., 2013) and is modelled as such. Determining whether the precise conformation of drug binding is influenced by the polymeric state of tubulin is an important question, but will only be possible once cryo-EM MT reconstructions are determined at well below 3Å resolution. In the currently modelled conformation, the macrocycle of epothilone is connected to β-tub-H7, the tip of S9-S10 and the M-loop, while the epothilone side-chain is directed towards the helical turn within the M-loop. The conformation of the binding pocket in Sp_tub is not detectably different in the presence of epothilone compared to its absence (Figure 3B).

**Figure 3.**
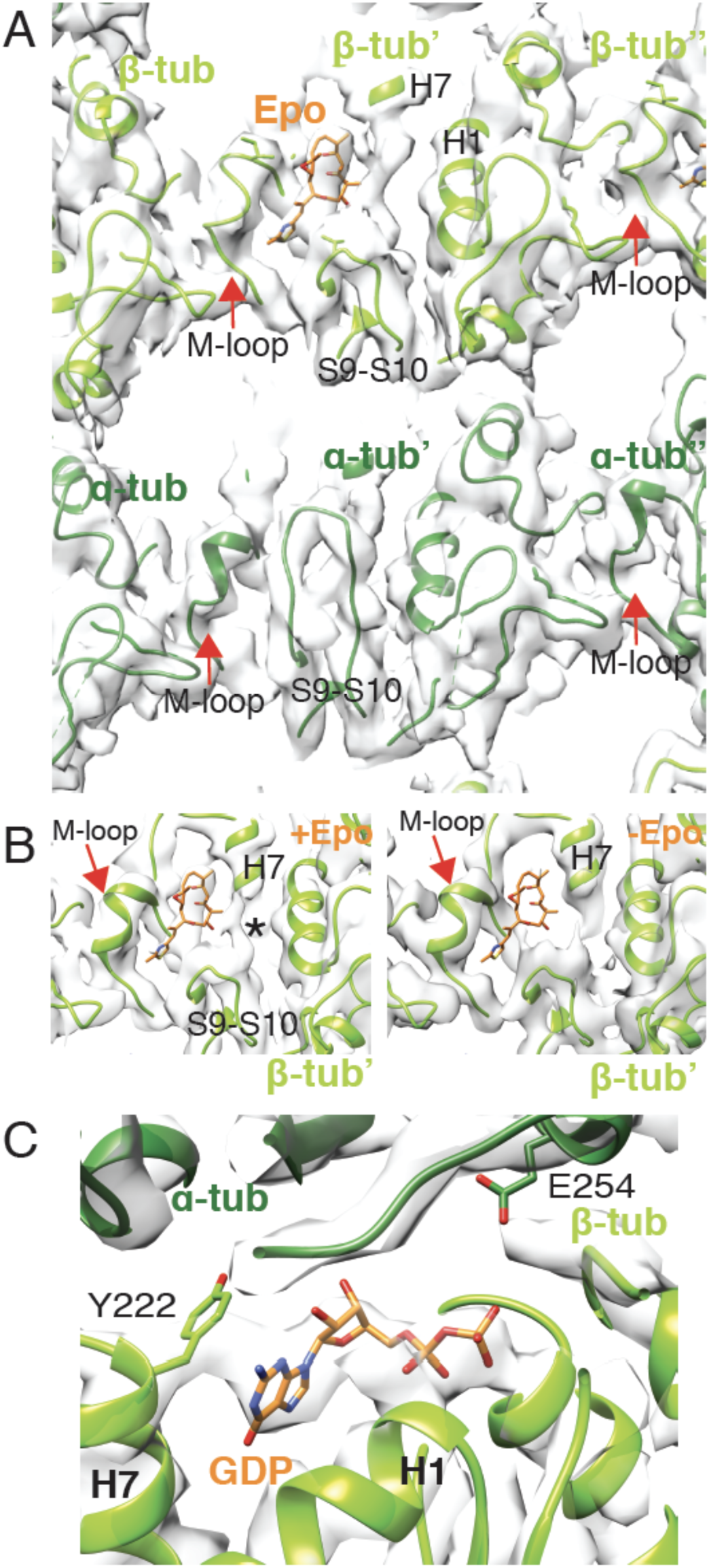
Epothilone binds at the β-tubulin taxane site of Sp_tub MTs. (A) Inter-PF lateral contacts viewed from the MT lumen, highlighting key secondary structure features and bound epothilone (Epo, orange); (B) Comparison of the taxane binding pocket in β-tubulin where density corresponding to epothilone is visible (left) with our previous structure of Sp_tub MTs without epothilone (right) (von Loeffelholz et al., 2017), showing the absence of density corresponding to the drug in a structure determined at equivalent resolution; an epothilone molecule is docked for comparison but lies outside this cryo-EM density; ^⋆^ in the left panel indicates unassigned density that may reflect mobility of the drug in the pocket; (C) Sp_tub β-tubulin E-site with β-tubulin in light green ribbon, α-tubulin in dark green ribbon, showing density consistent with bound GDP (in sticks).

Density in the nucleotide site of β-tubulin is consistent with bound GDP (Figure 3C). GTP hydrolysis and phosphate release likely occurs during sample preparation, as we observed previously with MT stabilization by binding protein Mal3 in the absence of epothilone (von Loeffelholz et al., 2017). Without addition of epothilone, these Sp_tub MTs depolymerize, suggesting that epothilone stabilizes Sp_tub MTs *in vitro*.

Mam_tub MTs are stabilized by paclitaxel, which binds in the same pocket on β-tubulin as epothilone but does not stabilize yeast (Sc_tub) MTs (Barnes et al., 1992). As an important chemotherapy drug, paclitaxel Mam_tub MT stabilization has been investigated extensively, although aspects of its stabilization mechanism remain under discussion (Alushin et al., 2014; Kellogg et al., 2017). Several recent studies have shown that the fundamental mechanochemistry of yeast tubulins is different compared to mammalian tubulins: specifically, the GTPase-dependent MT lattice compaction that is well-described in mammalian tubulin (Alushin et al., 2014; Hyman et al., 1995; Vale et al., 1994) either does not occur (von Loeffelholz et al., 2017) or is altered in yeast tubulins (Howes et al., 2017). Therefore, epothilone stabilization of yeast MTs is unlikely to depend on effects related to MT lattice compaction/expansion, but rather on other facets of MT structure, for example the lateral contacts between PFs. Given that paclitaxel and epothilone bind to the same site on MTs, in the future it will be important to learn more about the mechanochemical differences between these MTs. This will shed light on whether stabilization by drug binding at this pocket occurs by the same mechanism in MTs polymerized from different sources of tubulin, and to understand more about the impact on the motors that use these MTs as tracks.

In summary, our study sheds new light on diversity of structure and interaction between apparently well-conserved binding partners in the cytoskeleton, and highlights the importance of studying cognate motor-track combinations to elucidate physiologically relevant properties.

## Acknowledgements

O.v.L. and C.A.M. were supported by Biotechnology and Biological Sciences Research Council (BB/L00190X/1), A.P. and C.A.M. by WCR (16-0037), and thank the Wellcome Trust (079605/Z/06/Z) and BBSRC (BB/L014211/1) for EM equipment, Dr Charles Sindelar (Yale University, USA) for reconstruction algorithms, and the Birkbeck EM community for helpful discussions. R.A.C. is supported by a Wellcome Trust Senior Investigator Award (grant number 103895/Z/14/Z).

## Author Contributions

O.v.L. and A.P. conducted the experiments; D.R.D. provided vital reagents; all the authors were involved in designing the experiments and writing the paper.

## Declaration of Interests

The authors declare no competing interests

**Table 1.** Refinement statistics and model geometry for the Cut7/Sp_tub model. Refinement statistics and model geometry calculated in Phenix (Adams et al., 2010) and Molprobity (Chen et al., 2010).

|  |  |
| --- | --- |
| Resolution used for refinement | 4.5 Å |
| Molprobit score | 1.97 |
| RMSD (bonds, Å) | 0.01 |
| RMSD (angles) | 0.98 |
| All atoms clashscore | 8.35 |
| Ramachandran outliers | 0 % |
| Ramachandran allowed | 8.98% |
| Ramachandran favoured | 91.02% |
| Rotamer outliers | 0.2 % |

## Experimental Procedures

### Protein expression, purification and preparation

A recombinant His_6_-tagged Cut7 monomeric construct, residues 67–432 lacking its N-terminal extension (Cut7MD) in a pET151D-TOPO vector (Invitrogen) was expressed in BL21^⋆^(DE3) *Escherichia coli* cells as previously described (Britto et al., 2016). In brief, cells were grown in LB medium, supplemented with 2% (w/vol) glucose with induction of protein expression by 0.5 mM isopropyl β-D-1-thiogalactopyranoside (IPTG) at 18 °C for 5 h. Cells were resuspended in lysis buffer (50 mM Tris-HCl pH 8.0, 400 mM NaCl, 1 mM MgCl_2_, 1 mM ATP, 5 mM 2-mercaptoethanol, 10% (vol/vol) glycerol, and EDTA-free Protease Inhibitor Cocktail (Roche), 50 uM PMSF) and lysed using a French press. His_6_-tagged Cut7MD was purified from the clarified cell supernatant using nickel affinity chromatography and the His_6_ tag was removed using TEV protease during overnight dialysis into 50 mM Tris-HCl pH 8.0, 400 mM NaCl, 1 mM MgCl_2_, 1 mM ATP, 5 mM 2-mercaptoethanol, 10% (vol/vol) glycerol). Immediately prior to use, Cut7MD was buffer exchanged into BrB25+ (25 mM Pipes-KOH pH 6.8, 30 mM NaCl, 0.5 mM EGTA, 5 mM MgCl_2_, 1 mM 2-mercaptoethanol, 5 mM AMPPNP) using a Vivaspin^®^ column (Sartorius).

### Sample preparation for cryo-EM

*S. pombe* MTs were assembled from tag-free, dual isoform purified endogenous tubulin (Drummond et al., 2011) in PEM buffer (100 mM PIPES-KOH, 1 mM MgSO_4_, 2 mM EGTA, adjusted to pH 6.9 with KOH) mixed 1:1 with MES polymerization buffer (100 mM MES pH 6.5, 1 mM MgCl_2_, 1 mM EGTA, 1 mM DTT). 30 μM tubulin was polymerized in the presence of 5 mM GTP together with 25 μM monomeric Mal3 (residues 1-143), expressed in *E. coli* and purified as previously described (von Loeffelholz et al., 2017), except that the N-terminal His_6_ purification tag was removed by TEV protease cleavage. Monomeric Mal3 was added to bias the MT population to 13 PF architecture during polymerization in order to facilitate subsequent structure determination. MTs were polymerized at 32°C for 1 hour. Epothilone B (in DMSO (Stratech UK)) at a final concentration of 50 μM was added in the final 15 minutes of polymerization.

6 μM Sp_tub MT was mixed with 100 μM Cut7MD-AMPPNP at room temperature and 4μl of the mixture was applied immediately onto glow-discharged Quantifoil R 2/2 holey carbon grids which were blotted and plunge frozen into liquid ethane using a Vitrobot IV (FEI) operating at room temperature and 100% humidity.

### Cryo-EM data collection, structure determination and modelling

Movies were collected manually on a 300 kV Tecnai G2 Polara (FEI) microscope equipped with a Quantum energy filter and K2 Summit direct electron detector (Gatan) in counting mode, recording a total of 606 movies with a total dose in each of 30 e^-^/Å^2^ fractioned into 50 frames at a pixel size of 1.39 Å/px. Initial frame alignment was performed using IMOD (Kremer et al., 1996). A second local alignment step was performed with Scipion using the optical flow method (de la Rosa-Trevin et al., 2016). In the final reconstruction only frames 2-21 were included resulting in a total dose of 12 e^-^/Å^2^.

12543 MT segments were selected in 908 Å^2^ boxes in Boxer (Ludtke et al., 1999) using the helix option and choosing an overlap with three tubulin dimers (240 Å) unique in each box. Of the 748 MTs that were initially boxed, 669 MTs with 13_3 architecture were selected. The final 3D-reconstruction contained 33007 segments, reboxed with one unique tubulin dimer per box, and was calculated using a semi-automated single particle approach for pseudohelical assemblies in SPIDER and FREALIGN (Sindelar and Downing, 2010). Using the MT Fourier transform layer lines, the average helical repeat distance for the Sp_tub heterodimer in the epothilone-stabilized MTs was 82.98+/- 0.01 Å (mean and s.d., n=748 split into 3 groups). This is a small but statistically significant difference compared to the previously calculated GTP-Sp_tub = 83.26 +/- 0.01 Å (mean and s.d., (p<0.0001 (t-test)) and Mal3-143+GTP-Sp_tub = 82.91 +/- 0.02 Å (mean and s.d., p < 0.006 (t-test)) (von Loeffelholz et al., 2017).

The final reconstruction was automatically B-factor sharpened in RELION with an automated calculated B-factor of -234 (Scheres, 2012). As expected from the substoichiometric concentrations present during sample preparation, no density for the Mal3 added during MT polymerization was present in the final reconstruction. The overall resolution of the masked reconstruction was 4.5 Å (0.143 FSC). Local resolution was calculated with the blocres programme implemented in BSoft (Cardone et al., 2013), and structures were visualized using Chimera (Pettersen et al., 2004).

Homology models for the *S. pombe* αβ-tubulin dimer and Cut7 motor domain were adjusted from previous depositions (PDB: 5MJS and 5M5I, respectively) using Chimera and Coot (Emsley et al., 2010). The structure of epothilone B was downloaded from the grade web server (http://grade.globalphasing.org) and fitted into the density according to the previously determined high-resolution structure of the tubulin-stathmin-TTL-Epothilone complex (Prota et al., 2013). The atomic model was real space refined in Phenix (Adams et al., 2010) using the EM-map filtered to 4.5 Å. To improve the model geometry phenix.reduce (Word et al., 1999) was also used.

## Steady state ATPase assay

*S. pombe* MTs were assembled and stabilized by addition of epothilone as above, except that monomeric Mal3 was excluded from the polymerization mix. Mam_tub MTs were polymerized for 1 hour at 37°C, as previously described (Atherton et al., 2014) using bovine tubulin (Cytoskeleton Inc, Denver, CO) and stabilized by the addition of 1mM paclitaxel (Calbiochem). Cut7MD ATPase activity was measured using an enzyme-coupled assay (Kreuzer and Jongeneel, 1983) in a buffer consisting of 50 mM Tris (pH 8.0), 50 mM NaCl, 1 mM MgCl_2_, 0.5 mM phosphoenolpyruvate, 0.25 mM NADH, ~10 U/ml pyruvate kinase and ~14 U/ml lactate dehydrogenase and 5 mM ATP (all reagents from Sigma). The reactions were started by the addition of Cut7MD at a final concentration of 1.5 μM. Activity was measured by the decrease in NADH absorbance at 340 nm for 10 min at 32°C in a SpectraMax Plus 384 Microplate Reader (Molecular Devices).

## Data deposition

The cryo-EM reconstruction that supports the findings of this study has been deposited in the Electron Microscopy Data Bank, accession number 3527. The docked coordinates reported in this paper have been deposited in the Protein Data Bank, www.pdb.org, accession number 5MLV.

## Supplemental Information

### Supplemental Figures

**Figure S1 related to Figure 1.**
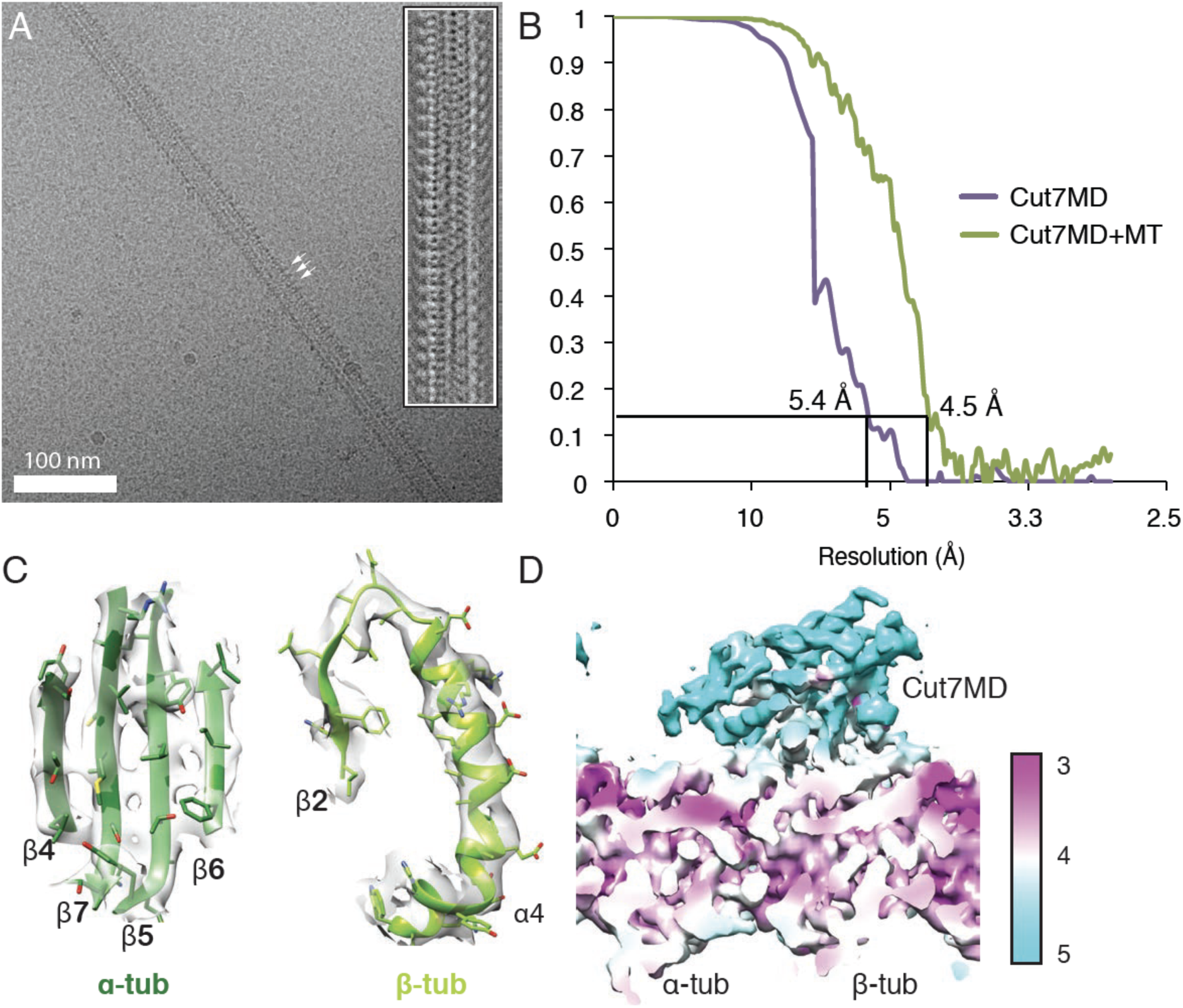
Visualization of Cut7MD binding to Sp_tub MTs and evaluation of the Cut7MD-Sp_tub MT cryo-EM reconstruction resolution. (A) Cryo-EM micrograph of Cut7MD-decorated Sp_tub MTs showing canonical binding of the motor domain every 8nm on the MT lattice (arrows); inset, MT segment average also shows clear Cut7MD binding every 8nm, as well as the characteristic moiré repeat of the Sp_tub 13PF MTs; (B) FSC curves for the overall reconstruction (green) estimated to be 4.5 Å by the 0.143 criterion and for the Cut7MD region specifically (purple); (C) Visualization of high resolution features in the Sp_tub density supports the near-atomic resolution quality of the reconstruction; (D) Depiction of local resolution estimate in the reconstruction using the *blocres* program implemented in Bsoft (Cardone et al., 2013) indicates the presence of a resolution gradient between the MT and kinesin.

**Figure S2 related to Figure 2.**
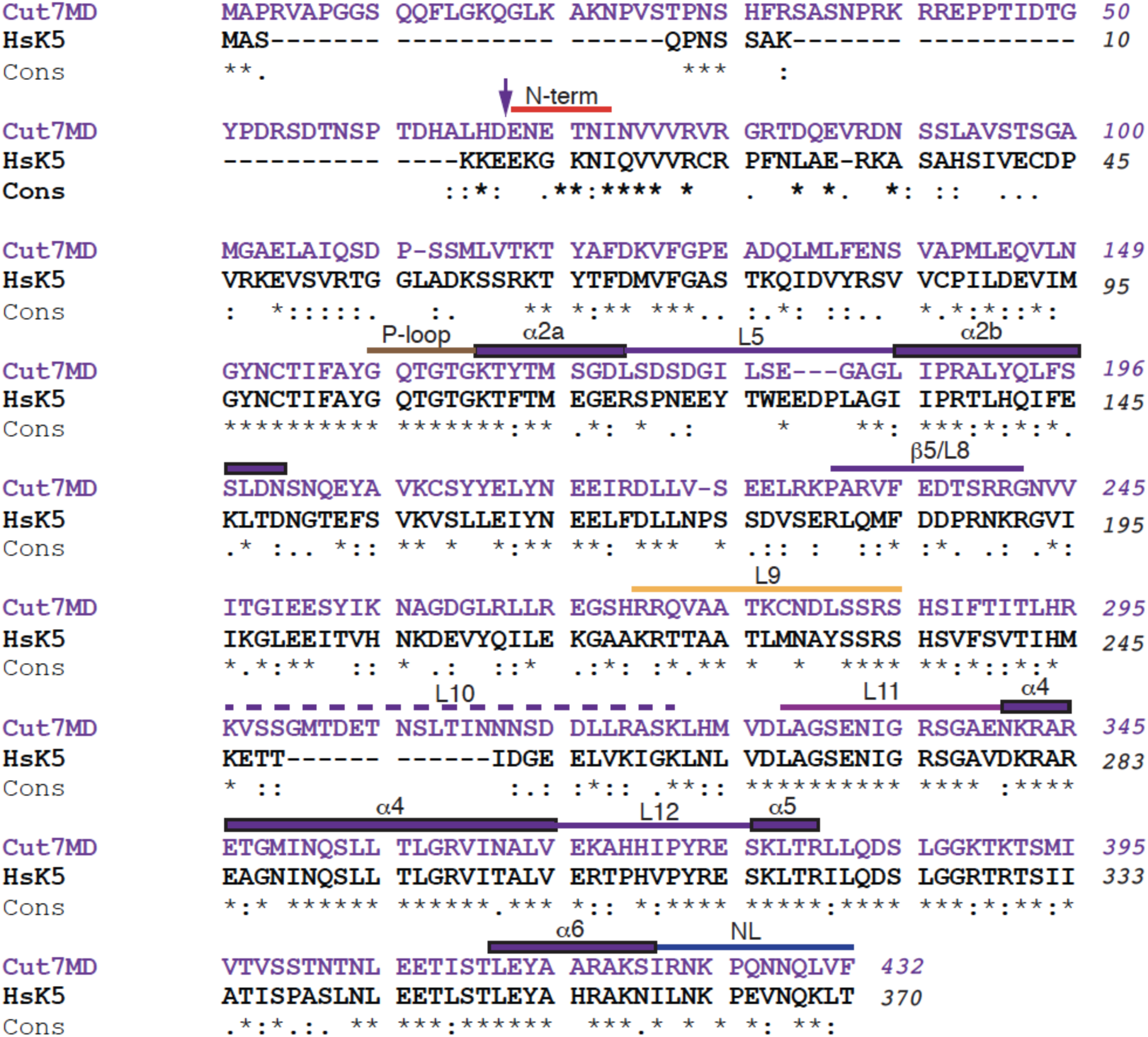
Sequence alignment of the Cut7 (Uniprot: P24339) and human Kif11 (Uniprot: P52732) motor domain sequences. Motor domain sequences were aligned using T-Coffee (Notredame et al., 2000) with sequence conservation depicted below the alignment. Secondary structure elements referenced in the text are annotated and the N-terminus of the Cut7MD construct used in this study is indicated with an arrow.

